# scEccDNAdb: an integrated single-cell eccDNA resource for human and mouse

**DOI:** 10.1101/2024.07.18.604058

**Authors:** Wenqing Wang, Xinyu Zhao, Tianyu Ma, Tengwei Zhong, Junnuo Zheng, Yi Yang, Yuanbiao Guo, Zhiyun Guo

## Abstract

Extrachromosomal circular DNA (eccDNA), an extrachromosomal circular structured DNA, is extensively found in eukaryotes. Exploring eccDNA at the single-cell level contributes to understanding heterogeneity, evolution, development, and specific functions within cells. Nevertheless, the high-throughput identification methods for single-cell eccDNA are complex, and currently mature and widely applicable technologies are lacking. Those factors have led to a scarcity of resources for studying eccDNA at the single-cell level. Therefore, using available single-cell whole-genome sequencing (WGS) data, we have constructed a comprehensive eccDNA database named scEccDNAdb (https://lcbb.swjtu.edu.cn/scEccDNAdb/). Presently, scEccDNAdb comprises 3,195,464 single-cell eccDNA entries from both disease/health human and mouse samples, which provides comprehensive annotations including oncogenes, typical enhancers, super-enhancers, CTCF binding sites, SNPs, chromatin accessibility, eQTLs, transcription factor binding sites, motifs, and SVs. Additionally, it provides nine online analysis and visualization tools, facilitating the generation of publication-quality figures for eccDNA analysis through the upload of customized files. Overall, scEccDNAdb represents the first comprehensive database known to us for exploring and analyzing single-cell eccDNA data in diverse cell types, tissues, and species.

**Graphical abstract:** 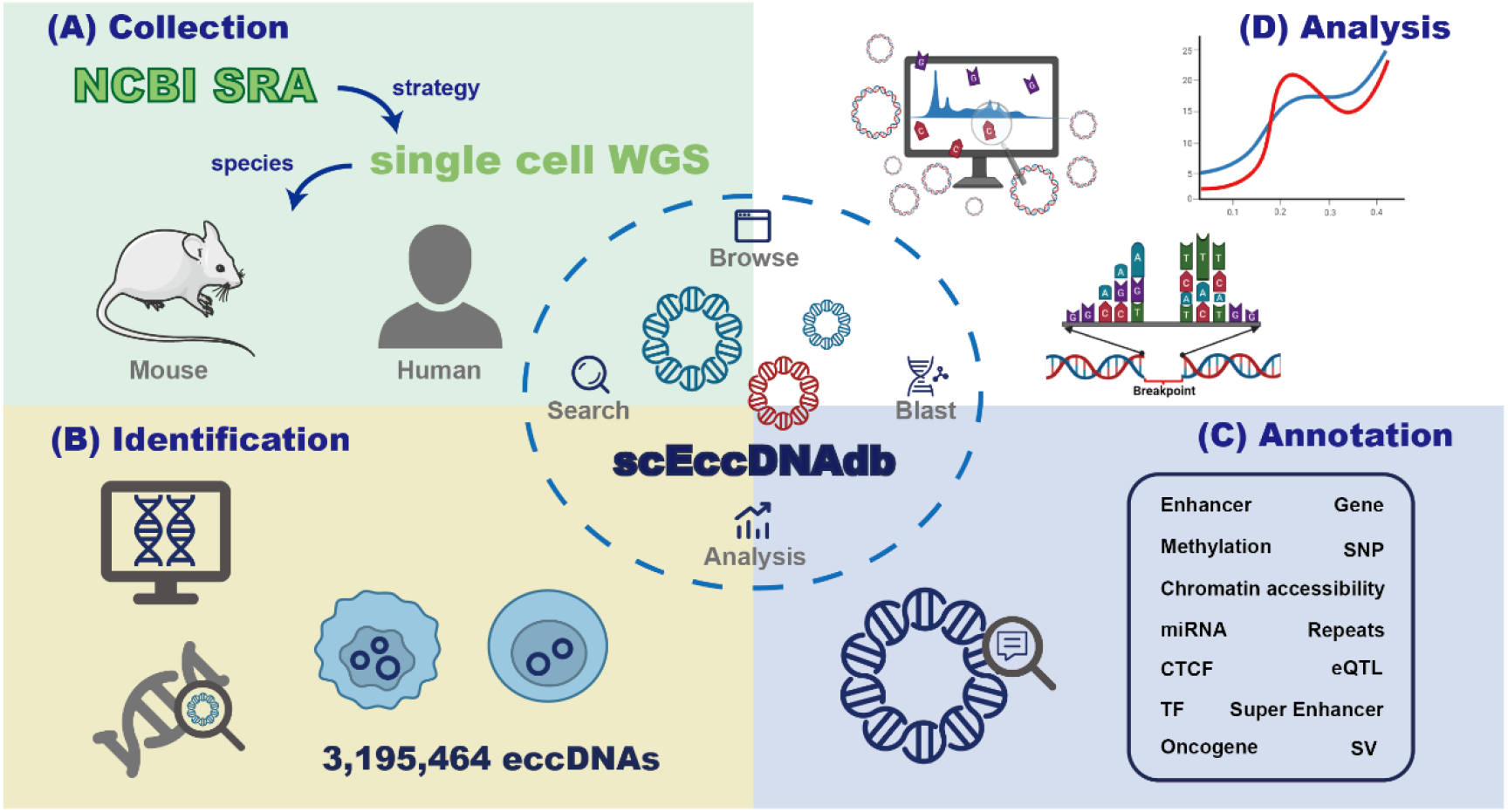

## Introduction

Extrachromosomal circular DNA (eccDNA), characterized by its circular structure outside the chromosomes, is widely present in eukaryotes and plays crucial roles in tumorigenesis and immune response, which can function as a biomarker (1). Due to the absence of centromeres, eccDNA is randomly distributed during mitosis in parental cells, resulting in heterogeneity in eccDNA quantity among daughter cells, and thereby causing cellular heterogeneity (2). In tumor cells, eccDNA-carrying oncogenes undergo high-copy amplification, consequently driving tumor progression and evolution (3). However, presently eccDNA researches predominantly focus on the bulk level, potentially equalizing the cellular diversity, which may cause the information loss in single-cell eccDNA. Moreover, some eccDNAs with low copy numbers or present in only a few cells cannot be detected using conventional bulk methods of eccDNA (4). In contrast, single-cell-based approaches can effectively overcome the above issues and provide deeper insights into the dynamic changes during the cell cycle, tracing its developmental process, cancer evolution (5,6), cellular functions, and regulation at the single-cell level (7-9).

Currently, single-cell eccDNA research primarily relies on sequencing technologies, such as SMOOTH-seq (9), scEC&T-seq (6), scCircle-seq (5), and scGTP-seq (10). However, challenges such as enrichment difficulty due to the circular nature of eccDNA and cost constraints, have interfered with the development of standardized high-throughput technologies for single-cell eccDNA identification. Currently, eccDNA identification methods based on whole-genome sequencing (WGS) data have been widely used. However, considering the extensive emergence of single-cell WGS data presently, these data are dispersed across various literature and datasets. Therefore, there is an urgent need to establish a repository for single-cell eccDNA research using these valuable single-cell WGS data.

To date, several eccDNA databases have been established, including CircleBase (11), eccDB (12), eccDNA Atlas (13), eccDNAdb (14), TeCD (15), and EccBase (16). Although these databases are very valuable, they are all bulk eccDNA databases, and there is currently no online resource for large-scale single-cell eccDNA collection and analysis. Here, we identified eccDNAs from 3,538 single-cell whole-genome sequencing datasets, and developed the first comprehensive single-cell eccDNA database as far as we known, named “scEccDNAdb”. We have developed a user-friendly web interface that enables users to explore detailed information and annotations related to eccDNA. Additionally, our database has multiple features such as search, blast, browse, and analysis, facilitating in-depth exploration of eccDNA data. Figure 1 illustrates the database construction process.

**Figure 1.**
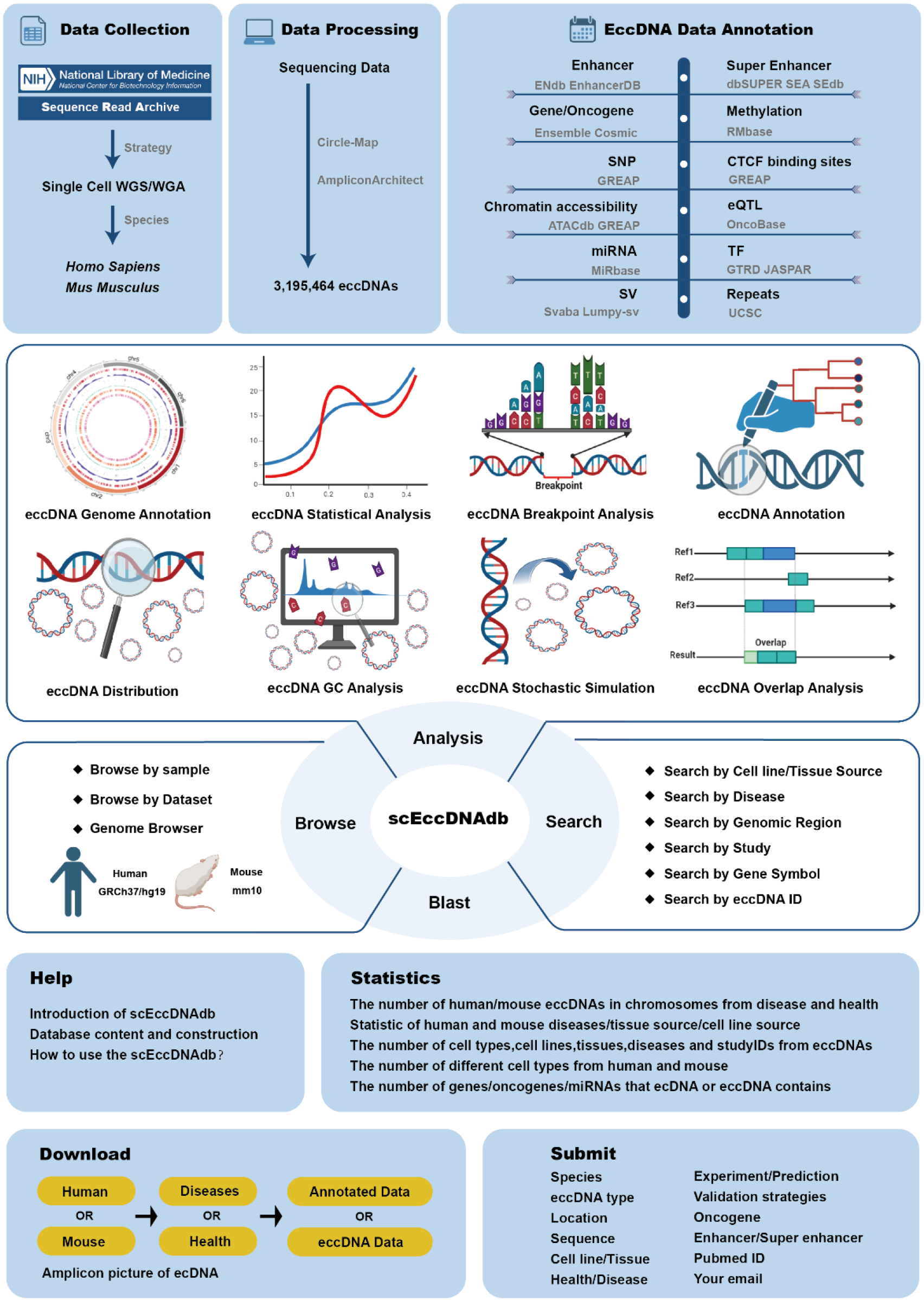
A schematic workflow of scEccDNAdb

## Materials and methods

### Data collection

In the NCBI-SRA database, single-cell whole-genome sequencing data for human and mouse were collected using the following three keywords: (a) ((((“single cell”[All Fields] OR “single-cell”[All Fields] OR “single cells”[All Fields] OR “single-cells”[All Fields]) AND WGA[Strategy]))); (b) ((((“single cell”[All Fields] OR “single-cell”[All Fields] OR “single cells”[All Fields] OR “single-cells”[All Fields]) AND WGS[Strategy]))); (c) “scWGS”. Duplicate data were removed post-retrieval, preserving paired-end data types, and then were manually corrected. A total of 2,163 disease and 1,027 health samples of single-cell human WGS data, and 79 disease and 269 health samples of single-cell mouse WGS data were obtained. Finally, the SRA toolkit was used for data download.

### Single-cell eccDNA identification

The quality control of WGS data was performed using Fastp (17) software with default parameters. Subsequently, BWA software was used to align reads to the reference genomes (human hg19, mouse mm10). Sorting and duplicate read filtering were performed using SAMtools (18) and Sambamba (19) software. EccDNA amplicons were identified using AmpliconArchitect (20) software based on the criteria of genome fragments >50 kb and copy numbers (CNs) ≥ 4.5. Finally, predicted amplicons were classified using AmpliconClassifier. Circle-Map (21) software was used for eccDNA identification, retaining results with “split read≥1” and “size <1kb”.

### Annotation and visualization

All eccDNA sequences collected by scEccDNAdb were converted to the default hg19 and mm10 genome version using the UCSC LiftOver tool (22). Regulatory elements located entirely within the eccDNA region were annotated as eccDNA regulatory elements. Detailed statistics are provided in Table 1. Circular plots of eccDNA were created using Circos (23). The UCSC browser was utilized for the visualization annotation of both the default existing and user-defined eccDNA regions, including histone modification signals, mutations, and sequence conservation.

**Table 1.**
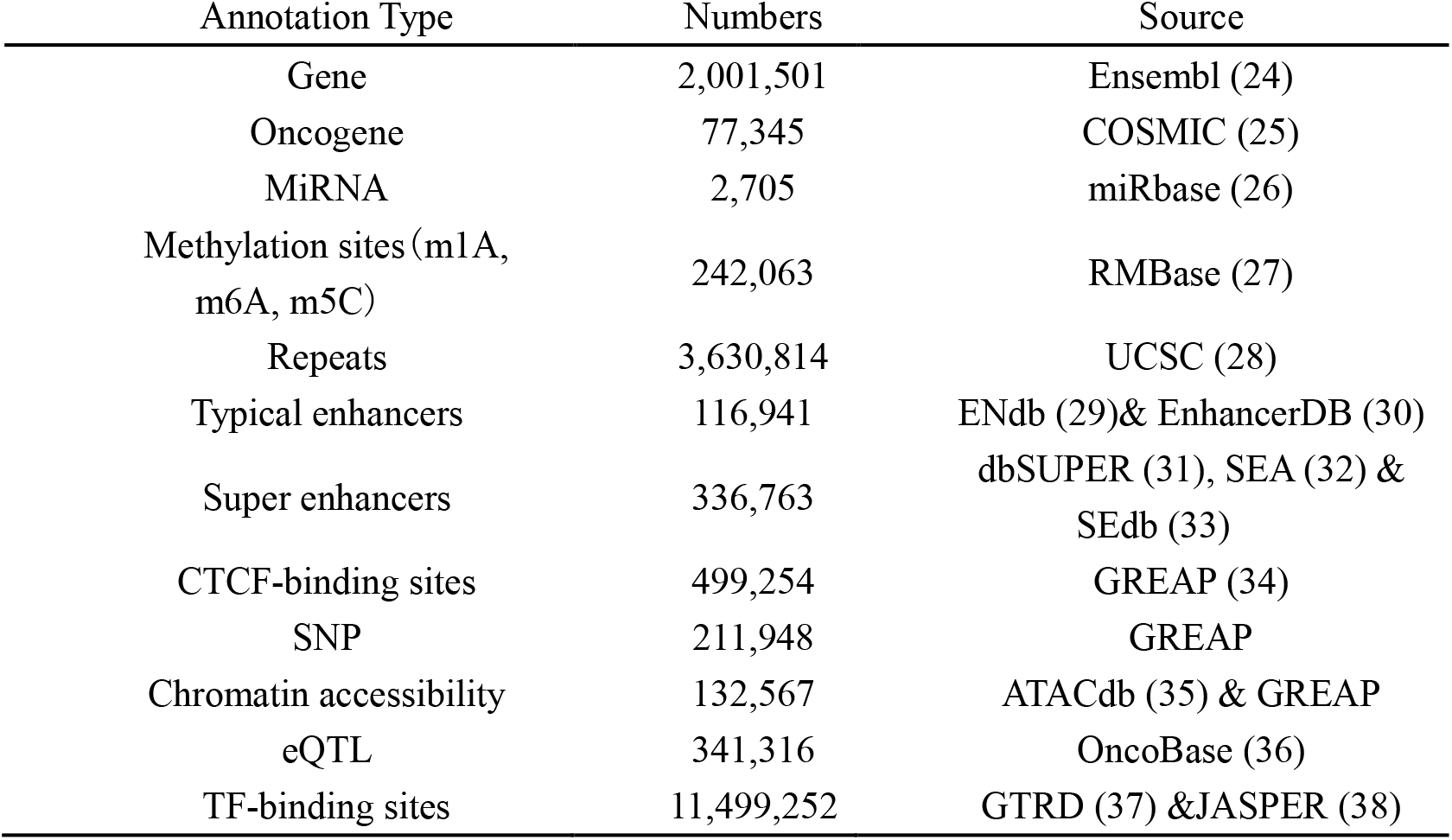
Annotation Data of eccDNAs in scEccDNAdb.

### Blast, structural variations, and motif enrichment analysis

BLAST was constructed using the ViroBLAST (39) online tool. MakeBlastdb was employed to create nucleic acid sequence libraries for human and mouse in scEccDNAdb. For human samples, sequences aligned to the reference genome were sorted and indexed using SAMtools, and then structural variant (SV) detection was performed for eccDNA regions using SvABA (40). For mouse samples, discordant paired-end alignments and split-read alignments were extracted from sequences aligned to the reference genome, and structural variant detection was performed using the lumpy-sv (41) software. Then, SQL was used to match the SV results above to each eccDNA of mouse. The Homer (v4.11)(42) software was used to perform motif enrichment analysis. The findMotifsGenome.pl command was used to detect motifs within each single-cell eccDNA genomic region with a size of 200bp.

### Analysis module development

The single-cell eccDNA analysis consists of eight modules, each developed and executed using R Shiny, with the system() function invoking Linux commands during analysis: 1) eccDNA Genome Annotation: A user-submitted function built in PHP enables users to input customized chromosome positions for annotation; 2) Statistical Analysis: Statistical graphs were generated using the R package ggplot2; 3) Breakpoint Analysis: The R package ggseqlogo (43) was used to analyze and plot sequence distribution patterns within the ± 10 bp region around the eccDNA breakpoints; 4) Annotation Analysis: The HOMER v4.11 script annotatePeaks.pl (hg19, with the -annStats parameter) was used to capture element types within specified eccDNA regions, with statistical graphs generated using the R package ggplot2; 5) Distribution Analysis: The R package Rideogram (44) was used to plot chromosome distribution of single-cell eccDNA; 6) GC Analysis: The computeMatrix scale-regions function in deeptools (45) v3.5.5 calculated signal density in specified regions, while the plotProfile function (with default parameters) generated signal plots of GC content in eccDNA regions. 7) Stochastic Simulation: The shuffle function in bedtools v2.31.0 (46) (with parameters -i, -excl, -g) was used to generate simulated regions similar to eccDNA regions submitted by the user. 8) Overlap Analysis: The intersect function in bedtools v2.31.0 (with parameters -a, -b, -wa, -wb, -f) captured eccDNA regions in scEccDNAdb that overlap with user-submitted eccDNA regions.

### System design and implementation

The scEccDNAdb website operates on a Nginx Web server (https://nginx.org/). The database was developed using MySQL 5.7.27 (http://www.mysql.com). PHP7.2.30 (http://www.php.net) was used for server-side scripting. The scEccDNAdb web interface was built using Bootstrap v3.3.7 (https://v3.bootcss.com) and JQuery v2.1.1 (http://jquery.com). ECharts (http://echarts.baidu.com) was used as a graphical visualization framework. We recommend using the latest versions of Firefox and Google Chrome for the best experience.

## Results

ScEccDNAdb consists of seven main sections: (i) BROWSE, (ii) SEARCH, (iii) ANALYSIS, (iv) BLAST, (v) DOWNLOAD, (vi) STATISTICS, and (vii) SUBMIT (Figure 2A). The detailed web interface and usage are as follows.

**Figure 2.**
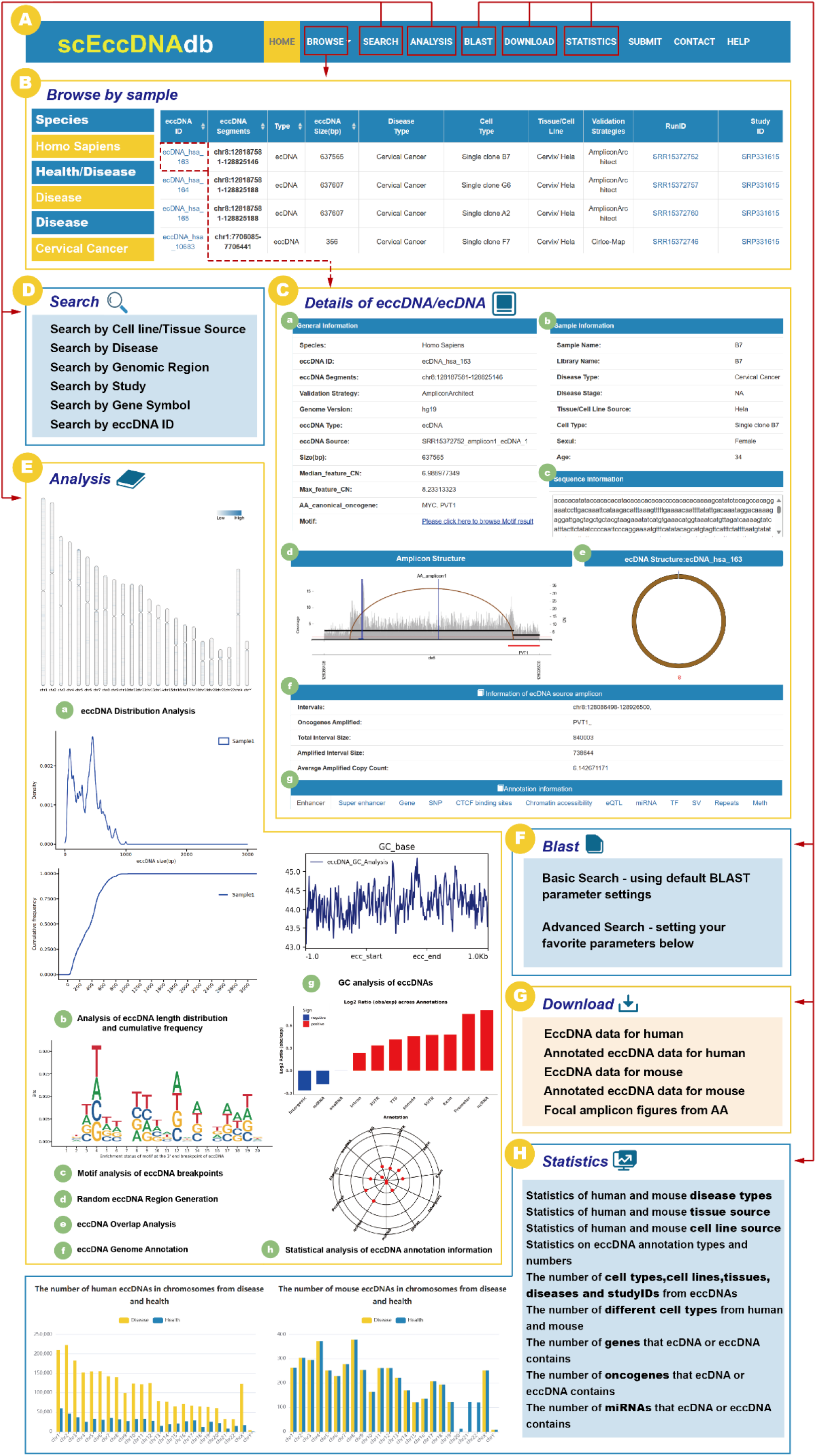
The web interface and usage of scEccDNAdb. (A) The navigation bar of scEccDNAdb. (B) Users can browse eccDNA data by sample and datasets. (C) “Details of eccDNA/ecDNA” includes: (a) General information, (b) Sample information, (c) Sequence information, (d) Amplicon structure, (e) Circular schematic of eccDNA, (f) Information of ecDNA source amplicon, and (g) Annotation information. (D) Users can query eccDNA data in six ways. (E) The eight analysis modules of scEccDNAdb. (F) Blast tool. (G) Data download page. (H)Statistics of scEccDNAdb.

### The browse and search module

The “Browse” page is divided into three menus: “Browse by sample,” “Browse by datasets,” and “Genome browser” (Figure 2B). “Browse by sample” enables users to explore eccDNA by species, disease type (Health/Disease), and cell type (tissue source/cell line source). “Browse by datasets” allows users to browse eccDNA by species, disease type, and study ID. Users can click on an eccDNA ID to access detailed information about that eccDNA (Figure 2C). The eccDNA detail information includes comprehensive coverage of eccDNA chromosome regions, general information, sample information, sequence, amplicon structure, circular schematic of eccDNA, amplification details, annotation information, etc. The “Genome Browser” is a visualization tool based on the UCSC genome browser, enabling users to visualize eccDNA genomic regions. Users can select a specific eccDNA from scEccDNAdb or enter a custom eccDNA genomic region to visualize histone modification signals, mutations, sequence conservation, and other relevant information within the eccDNA region.

In the SEARCH module, users can conduct queries on eccDNA data using six approaches: (i) “Search by tissue/cell line source” for searching eccDNAs through three steps, including “species”, “Disease/Health”, and “cell line”; (ii) “Search by Disease” for searching eccDNAs based on different disease types in humans or mouse; (iii) “Search by region” for searching chromosome locations of interest by users of all eccDNAs we collected; (iv) “Search by study” for searching eccDNAs from the studies used for eccDNA identification; (v) “Search by gene symbol” for searching the gene on eccDNA by name; (vi) “Search by eccDNA ID” (Figure 2D).

### EccDNA single-cell analysis module

The ANALYSIS module of scEccDNAdb includes the following eight parts (Figure 2E): (a) “eccDNA Distribution Analysis” generates publication-quality figures for eccDNA chromosomal distribution in a single or paired sample by uploading a customized file. (b) “eccDNA Statistical Analysis” generates publication-quality figures for length distribution and cumulative frequency of eccDNA in a single or paired sample by uploading a customized bed file. (c) “eccDNA Breakpoint Analysis” generates publication-quality sequence logos (motif enrichment) in the ±10 bp region around the eccDNA breakpoints by uploading a customized file. (d) “eccDNA Stochastic Simulation” generates random eccDNA regions of the same size as the user uploads a customized file of eccDNA regions. (e) “eccDNA Overlap Analysis” performs overlap analysis between customized eccDNA regions and eccDNA regions of scEccDNAdb. (f) “eccDNA Genome Annotation” performs annotation analysis on eccDNA regions included in scEccDNAdb or custom chromatin regions, including oncogenes/lncRNAs, typical enhancers, super-enhancers, CTCF binding sites, SNPs, chromatin accessibility, and eQTLs. Furthermore, we provide external links for eccDNA details, including GEPIA (47), GeneCard (48), and String databases (49). (g) “eccDNA GC Analysis” generates publication-quality GC enrichment figures of eccDNAs by uploading a customized file. (h) “eccDNA Annotation Analysis” observes the enrichment and quantity distribution of annotated elements of single-cell eccDNA for a specific time period.

### BLAST

To assist users in identifying regions of local similarity between query sequences and eccDNA sequences in scEccDNAdb, we have incorporated the BLAST tool on the webpage. Users can upload or paste DNA sequences in FASTA format and select different datasets for the BLAST search. Furthermore, the user can conduct a more refined search by adjusting filtering parameters (Figure 2F).

### Download, submit and statistics

Users can download all eccDNA, annotation, and analysis data from scEccDNAdb (Figure 2G) and share their research data (e.g., location, sequence) through the submission interface. It should be noted that data submitted by users will not be made available online immediately until a rigorous manual verification process is completed. On the statistics page, eccDNA is primarily classified and counted according to chromosome, tissue, cell line, disease, etc. (Figure 2H).

### A case study

Taking single-cell eccDNA of cervical cancer as an example, the detailed application of scEccDNAdb is as follows. In the “browse by sample” interface, users can access a list of data containing all eccDNA samples from single cells of cervical cancer by clicking on the “Cervical Cancer” label. In the search interface, using the “Search eccDNA by Disease” strategy, users can select “Homo sapiens” as the species and “Cervical Cancer” as the disease type in Steps 1 and 2, respectively, to retrieve data on 4,626 eccDNAs from cervical cancer samples. The search results page provides information including eccDNA ID, eccDNA segments, eccDNA size (bp), disease type, cell type, tissue/cell type, validation strategies, run ID, and study ID for each eccDNA. Furthermore, users can refine the results list using the “Search” box located at the top left corner.

Taking “ecDNA_hsa_163” from the results list as an example, users can click on the eccDNA ID to gain detailed insights into the specific eccDNA. The “General information” in the “Detail” interface reveals that this eccDNA spans the region chr8:128187581-128825146, sourced from single-cell clone sample “B7” of the “Hela” cell line, with a size of 637,565 bp. The “Sequence information” presents the nucleotide arrangement of this eccDNA region. The “Median_feature_CN” is “6.988977349”, and the “Max_feature_CN” is “8.23313323”. Notably, the annotation identifies the oncogenes “MYC” and “PVT1” within ecDNA_hsa_163, both known to play crucial roles in tumorigenesis. Previous studies have shown that the PVT1 promoter is ectopically fused with MYC and duplicated in ecDNA, resulting in potent MYC expression mediated through enhancer hijacking (50). The ecDNA structure diagram illustrates the circular structure of ecDNA_hsa_163. The “Amplicon structure” delineates the amplification structure of the eccDNA source, while the “Information of ecDNA source amplicon” section illustrates pertinent details such as the interval located at chr8:128086498-128926500, amplified oncogenes including PVT1, and total interval size of 840,003bp. Finally, the “annotation information” encompasses annotations on the ecDNA_hsa_163, including 55 enhancers, 40 super-enhancers, nine genes (including the oncogenes MYC and PVT1), 3,534 CTCF binding sites, 5,117 chromatin accessibility regions, 31 eQTLs, two miRNAs, 494 TFs, 1,441 SVs, 1,436 repeats, and 80 methylation regions. In conclusion, scEccDNAdb is crucial for advancing research into the potential functions and regulatory roles of eccDNAs in diseases.

## Discussion and future development

Here we have collected available publicly single-cell WGS data to identify eccDNA and developed the scEccDNAdb database. scEccDNAdb is a unique comprehensive repository for single-cell eccDNA, providing extensive datasets derived from human and mouse single-cell samples, including sample metadata, genomic sequences, and thorough annotations. Moreover, scEccDNAdb provides online analysis tools and visualization tools to help users explore and deepen understanding of eccDNA biogenesis and regulatory mechanisms. However, there are also some challenges in identifying and analyzing eccDNA. For example, single-cell WGS data are often low-coverage WGS, effectively affecting eccDNA identification. Additionally, the lack of omics data correlating with cell types and clinical information brings challenges for a comprehensive analysis of single-cell eccDNA.

In the future, with the development of single-cell technologies, more and more omics data will be generated, contributing to a deeper understanding of eccDNA. We will continually collect the latest single-cell datasets and add these datasets to scEccDNAdb. Furthermore, we plan to integrate more relevant eccDNA analysis tools for conjoint analysis of eccDNA, including eccDNA-pipe (51), CReSIL (52), Circlehunter (53), DeepCircle (54), and ATACAmp (55), etc. We will also develop new data analysis features to enhance the database’s interactivity, providing users with an enhanced experience. By consistently using and refining scEccDNAdb, our goal is to deepen our understanding of eccDNA single-cell heterogeneity especially its roles in intratumoral heterogeneity and cancer evolution.

## Data availability

The scEccDNAdb is publicly accessible at https://lcbb.swjtu.edu.cn/scEccDNAdb/. Users can directly download the data resources without registration or login.

## Acknowledgments

We thank the Informatization and Network Management Office of Southwest Jiaotong University for providing cloud-based technical support.

## References

1. Zhao, X., Zhao, H., Liu, Y. and Guo, Z. (2023) Methods, bioinformatics tools and databases in ecDNA research: An overview. Comput Biol Med, 167, 107680.

2. Wu, S., Bafna, V., Chang, H.Y. and Mischel, P.S. (2022) Extrachromosomal DNA: An Emerging Hallmark in Human Cancer. Annu Rev Pathol, 17, 367–386.

3. Turner, K.M., Deshpande, V., Beyter, D., Koga, T., Rusert, J., Lee, C., Li, B., Arden, K., Ren, B., Nathanson, D.A. et al. (2017) Extrachromosomal oncogene amplification drives tumour evolution and genetic heterogeneity. Nature, 543, 122–125.

4. Jiang, R., Yang, M., Zhang, S. and Huang, M. (2023) Advances in sequencing-based studies of microDNA and ecDNA: Databases, identification methods, and integration with single-cell analysis. Comput Struct Biotechnol J, 21, 3073–3080.

5. Chen, J.P., Diekmann, C., Wu, H., Chen, C., Della Chiara, G., Berrino, E., Georgiadis, K.L., Bouwman, B.A.M., Virdi, M., Harbers, L. et al. (2024) scCircle-seq unveils the diversity and complexity of extrachromosomal circular DNAs in single cells. Nat Commun, 15, 1768.

6. Chamorro Gonzalez, R., Conrad, T., Stober, M.C., Xu, R., Giurgiu, M., Rodriguez-Fos, E., Kasack, K., Bruckner, L., van Leen, E., Helmsauer, K. et al. (2023) Parallel sequencing of extrachromosomal circular DNAs and transcriptomes in single cancer cells. Nat Genet, 55, 880–890.

7. Kang, J., Dai, Y., Li, J., Fan, H. and Zhao, Z. (2023) Investigating cellular heterogeneity at the single-cell level by the flexible and mobile extrachromosomal circular DNA. Comput Struct Biotechnol J, 21, 1115–1121.

8. Wen, L. and Tang, F. (2022) Recent advances in single-cell sequencing technologies. Precis Clin Med, 5, pbac002.

9. Fan, X., Yang, C., Li, W., Bai, X., Zhou, X., Xie, H., Wen, L. and Tang, F. (2021) SMOOTH-seq: single-cell genome sequencing of human cells on a third-generation sequencing platform. Genome Biol, 22, 195.

10. Chang, L., Deng, E., Wang, J., Zhou, W., Ao, J., Liu, R., Su, D. and Fan, X. (2023) Single-cell third-generation sequencing-based multi-omics uncovers gene expression changes governed by ecDNA and structural variants in cancer cells. Clin Transl Med, 13, e1351.

11. Zhao, X., Shi, L., Ruan, S., Bi, W., Chen, Y., Chen, L., Liu, Y., Li, M., Qiao, J. and Mao, F. (2022) CircleBase: an integrated resource and analysis platform for human eccDNAs. Nucleic Acids Res, 50, D72–D82.

12. Yang, M., Qiu, B., He, G.Y., Zhou, J.Y., Yu, H.J., Zhang, Y.Y., Li, Y.S., Li, T.S., Guo, J.C., Li, X.C. et al. (2023) eccDB: a comprehensive repository for eccDNA-mediated chromatin contacts in multi-species. Bioinformatics, 39.

13. Zhong, T., Wang, W., Liu, H., Zeng, M., Zhao, X. and Guo, Z. (2023) eccDNA Atlas: a comprehensive resource of eccDNA catalog. Brief Bioinform, 24.

14. Peng, L., Zhou, N., Zhang, C.Y., Li, G.C. and Yuan, X.Q. (2022) eccDNAdb: a database of extrachromosomal circular DNA profiles in human cancers. Oncogene, 41, 2696–2705.

15. Guo, J., Zhang, Z., Li, Q., Chang, X. and Liu, X. (2023) TeCD: The eccDNA Collection Database for extrachromosomal circular DNA. BMC Genomics, 24, 47.

16. Sun, H., Lu, X. and Zou, L. (2023) EccBase: A high-quality database for exploration and characterization of extrachromosomal circular DNAs in cancer. Comput Struct Biotechnol J, 21, 2591–2601.

17. Chen, S., Zhou, Y., Chen, Y. and Gu, J. (2018) fastp: an ultra-fast all-in-one FASTQ preprocessor. Bioinformatics, 34, i884–i890.

18. Danecek, P., Bonfield, J.K., Liddle, J., Marshall, J., Ohan, V., Pollard, M.O., Whitwham, A., Keane, T., McCarthy, S.A., Davies, R.M. et al. (2021) Twelve years of SAMtools and BCFtools. Gigascience, 10.

19. Tarasov, A., Vilella, A.J., Cuppen, E., Nijman, I.J. and Prins, P. (2015) Sambamba: fast processing of NGS alignment formats. Bioinformatics, 31, 2032–2034.

20. Deshpande, V., Luebeck, J., Nguyen, N.D., Bakhtiari, M., Turner, K.M., Schwab, R., Carter, H., Mischel, P.S. and Bafna, V. (2019) Exploring the landscape of focal amplifications in cancer using AmpliconArchitect. Nat Commun, 10, 392.

21. Prada-Luengo, I., Krogh, A., Maretty, L. and Regenberg, B. (2019) Sensitive detection of circular DNAs at single-nucleotide resolution using guided realignment of partially aligned reads. BMC Bioinformatics, 20, 663.

22. Rosenbloom, K.R., Armstrong, J., Barber, G.P., Casper, J., Clawson, H., Diekhans, M., Dreszer, T.R., Fujita, P.A., Guruvadoo, L., Haeussler, M. et al. (2015) The UCSC Genome Browser database: 2015 update. Nucleic Acids Res, 43, D670–681.

23. Krzywinski, M., Schein, J., Birol, I., Connors, J., Gascoyne, R., Horsman, D., Jones, S.J. and Marra, M.A. (2009) Circos: an information aesthetic for comparative genomics. Genome Res, 19, 1639–1645.

24. Harrison, P.W., Amode, M.R., Austine-Orimoloye, O., Azov, A.G., Barba, M., Barnes, I., Becker, A., Bennett, R., Berry, A., Bhai, J. et al. (2024) Ensembl 2024. Nucleic Acids Res, 52, D891–D899.

25. Tate, J.G., Bamford, S., Jubb, H.C., Sondka, Z., Beare, D.M., Bindal, N., Boutselakis, H., Cole, C.G., Creatore, C., Dawson, E. et al. (2019) COSMIC: the Catalogue Of Somatic Mutations In Cancer. Nucleic Acids Res, 47, D941–D947.

26. Kozomara, A., Birgaoanu, M. and Griffiths-Jones, S. (2019) miRBase: from microRNA sequences to function. Nucleic Acids Res, 47, D155–D162.

27. Sun, W.J., Li, J.H., Liu, S., Wu, J., Zhou, H., Qu, L.H. and Yang, J.H. (2016) RMBase: a resource for decoding the landscape of RNA modifications from high-throughput sequencing data. Nucleic Acids Res, 44, D259–265.

28. Nassar, L.R., Barber, G.P., Benet-Pages, A., Casper, J., Clawson, H., Diekhans, M., Fischer, C., Gonzalez, J.N., Hinrichs, A.S., Lee, B.T. et al. (2023) The UCSC Genome Browser database: 2023 update. Nucleic Acids Res, 51, D1188–D1195.

29. Bai, X., Shi, S., Ai, B., Jiang, Y., Liu, Y., Han, X., Xu, M., Pan, Q., Wang, F., Wang, Q. et al. (2020) ENdb: a manually curated database of experimentally supported enhancers for human and mouse. Nucleic Acids Res, 48, D51–D57.

30. Kang, R., Zhang, Y., Huang, Q., Meng, J., Ding, R., Chang, Y., Xiong, L. and Guo, Z. (2019) EnhancerDB: a resource of transcriptional regulation in the context of enhancers. Database (Oxford), 2019.

31. Khan, A. and Zhang, X. (2016) dbSUPER: a database of super-enhancers in mouse and human genome. Nucleic Acids Res, 44, D164–171.

32. Chen, C., Zhou, D., Gu, Y., Wang, C., Zhang, M., Lin, X., Xing, J., Wang, H. and Zhang, Y. (2020) SEA version 3.0: a comprehensive extension and update of the Super-Enhancer archive. Nucleic Acids Res, 48, D198–D203.

33. Jiang, Y., Qian, F., Bai, X., Liu, Y., Wang, Q., Ai, B., Han, X., Shi, S., Zhang, J., Li, X. et al. (2019) SEdb: a comprehensive human super-enhancer database. Nucleic Acids Res, 47, D235–D243.

34. Yang, Y., Qian, F., Li, X., Li, Y., Zhou, L., Wang, Q., Zhou, X., Zhang, J., Song, C., Yu, Z. et al. (2022) GREAP: a comprehensive enrichment analysis software for human genomic regions. Brief Bioinform, 23.

35. Wang, F., Bai, X., Wang, Y., Jiang, Y., Ai, B., Zhang, Y., Liu, Y., Xu, M., Wang, Q., Han, X. et al. (2021) ATACdb: a comprehensive human chromatin accessibility database. Nucleic Acids Res, 49, D55–D64.

36. Li, X., Shi, L., Wang, Y., Zhong, J., Zhao, X., Teng, H., Shi, X., Yang, H., Ruan, S., Li, M. et al. (2019) OncoBase: a platform for decoding regulatory somatic mutations in human cancers. Nucleic Acids Res, 47, D1044–D1055.

37. Kolmykov, S., Yevshin, I., Kulyashov, M., Sharipov, R., Kondrakhin, Y., Makeev, V.J., Kulakovskiy, I.V., Kel, A. and Kolpakov, F. (2021) GTRD: an integrated view of transcription regulation. Nucleic Acids Res, 49, D104–D111.

38. Rauluseviciute, I., Riudavets-Puig, R., Blanc-Mathieu, R., Castro-Mondragon, J.A., Ferenc, K., Kumar, V., Lemma, R.B., Lucas, J., Cheneby, J., Baranasic, D. et al. (2024) JASPAR 2024: 20th anniversary of the open-access database of transcription factor binding profiles. Nucleic Acids Res, 52, D174–D182.

39. Deng, W., Nickle, D.C., Learn, G.H., Maust, B. and Mullins, J.I. (2007) ViroBLAST: a stand-alone BLAST web server for flexible queries of multiple databases and user’s datasets. Bioinformatics, 23, 2334–2336.

40. Wala, J.A., Bandopadhayay, P., Greenwald, N.F., O’Rourke, R., Sharpe, T., Stewart, C., Schumacher, S., Li, Y., Weischenfeldt, J., Yao, X. et al. (2018) SvABA: genome-wide detection of structural variants and indels by local assembly. Genome Res, 28, 581–591.

41. Layer, R.M., Chiang, C., Quinlan, A.R. and Hall, I.M. (2014) LUMPY: a probabilistic framework for structural variant discovery. Genome Biol, 15, R84.

42. Heinz, S., Benner, C., Spann, N., Bertolino, E., Lin, Y.C., Laslo, P., Cheng, J.X., Murre, C., Singh, H. and Glass, C.K. (2010) Simple combinations of lineage-determining transcription factors prime cis-regulatory elements required for macrophage and B cell identities. Mol Cell, 38, 576–589.

43. Wagih, O. (2017) ggseqlogo: a versatile R package for drawing sequence logos. Bioinformatics, 33, 3645–3647.

44. Hao, Z., Lv, D., Ge, Y., Shi, J., Weijers, D., Yu, G. and Chen, J. (2020) RIdeogram: drawing SVG graphics to visualize and map genome-wide data on the idiograms. PeerJ Comput Sci, 6, e251.

45. Ramirez, F., Dundar, F., Diehl, S., Gruning, B.A. and Manke, T. (2014) deepTools: a flexible platform for exploring deep-sequencing data. Nucleic Acids Res, 42, W187–191.

46. Quinlan, A.R. and Hall, I.M. (2010) BEDTools: a flexible suite of utilities for comparing genomic features. Bioinformatics, 26, 841–842.

47. Tang, Z., Li, C., Kang, B., Gao, G., Li, C. and Zhang, Z. (2017) GEPIA: a web server for cancer and normal gene expression profiling and interactive analyses. Nucleic Acids Res, 45, W98–W102.

48. Stelzer, G., Rosen, N., Plaschkes, I., Zimmerman, S., Twik, M., Fishilevich, S., Stein, T.I., Nudel, R., Lieder, I., Mazor, Y. et al. (2016) The GeneCards Suite: From Gene Data Mining to Disease Genome Sequence Analyses. Curr Protoc Bioinformatics, 54, 1 30 31-31 30 33.

49. Szklarczyk, D., Kirsch, R., Koutrouli, M., Nastou, K., Mehryary, F., Hachilif, R., Gable, A.L., Fang, T., Doncheva, N.T., Pyysalo, S. et al. (2023) The STRING database in 2023: protein-protein association networks and functional enrichment analyses for any sequenced genome of interest. Nucleic Acids Res, 51, D638–D646.

50. Hung, K.L., Yost, K.E., Xie, L., Shi, Q., Helmsauer, K., Luebeck, J., Schopflin, R., Lange, J.T., Chamorro Gonzalez, R., Weiser, N.E. et al. (2021) ecDNA hubs drive cooperative intermolecular oncogene expression. Nature, 600, 731–736.

51. Fang, M., Fang, J., Luo, S., Liu, K., Yu, Q., Yang, J., Zhou, Y., Li, Z., Sun, R., Guo, C. et al. (2024) eccDNA-pipe: an integrated pipeline for identification, analysis and visualization of extrachromosomal circular DNA from high-throughput sequencing data. Brief Bioinform, 25.

52. Wanchai, V., Jenjaroenpun, P., Leangapichart, T., Arrey, G., Burnham, C.M., Tummler, M.C., Delgado-Calle, J., Regenberg, B. and Nookaew, I. (2022) CReSIL: accurate identification of extrachromosomal circular DNA from long-read sequences. Brief Bioinform, 23.

53. Yang, M., Zhang, S., Jiang, R., Chen, S. and Huang, M. (2023) Circlehunter: a tool to identify extrachromosomal circular DNA from ATAC-Seq data. Oncogenesis, 12, 28.

54. Chang, K.L., Chen, J.H., Lin, T.C., Leu, J.Y., Kao, C.F., Wong, J.Y. and Tsai, H.K. (2023) Short human eccDNAs are predictable from sequences. Brief Bioinform, 24.

55. Cheng, H., Ma, W., Wang, K., Chu, H., Bao, G., Liao, Y., Yuan, Y., Gou, Y., Dong, L., Yang, J. et al. (2023) ATACAmp: a tool for detecting ecDNA/HSRs from bulk and single-cell ATAC-seq data. BMC Genomics, 24, 678.

